# Territory-level temperature influences breeding phenology and reproductive output in three forest passerine birds

**DOI:** 10.1101/2021.01.31.429009

**Authors:** Jack D Shutt, Sophie C Bell, Fraser Bell, Joan Castello, Myriam El Harouchi, Malcolm D Burgess

## Abstract

Temperature plays an important role in determining breeding phenology of temperate birds, with higher spring temperatures associated with earlier breeding. However, the effect of localised territory-scale temperature variations is poorly understood, with relationships between temperature and breeding phenology mostly studied using coarse-grained climatic indices. Here, we interpolate spring temperatures recorded at 150 m^2^ grid intersections encompassing 417 ha of forest to examine the influence of territory-scale temperature, and its interaction with mean annual temperature, on territory selection, breeding phenology, clutch size and fledgling success for three co-occurring single-brooded passerine birds using data from 672 nests over four years. All species exhibited significant trends in reproductive traits associated with territory-scale temperature. Pied flycatchers *Ficedula hypoleuca* settled in cooler territories first, where they raised more fledglings. Blue tits *Cyanistes caeruleus* laid larger clutches in warmer territories in warm years and always laid earlier at warmer territories irrespective of annual temperature variation. Contrastingly, pied flycatcher and wood warbler *Phylloscopus sibilatrix* breeding phenology was earlier at warmer territories in cool years and cooler territories in warm years, with wood warbler clutch size responding similarly to this interaction. Greater previous breeding experience and increased higher rates of historical territory occupancy (territory quality) also predicted earlier breeding phenology and higher fledging success for pied flycatchers. We suggest that the migratory pied flycatcher and wood warbler are best synchronised with their prey availability in cooler years at a local population level while resident blue tits match local phenology across all years, which is potentially advantageous under warmer predicted climate change scenarios. We conclude that temperature at the territory scale can be an important driver of settlement and breeding phenology and influence reproductive traits.

## Introduction

Temperature plays a key role in determining the timing of breeding of temperate zone birds (Visser, Holleman & Caro 2009). In warmer years, and at generally warmer locations, reproduction occurs earlier, and under recent climate change average breeding phenology has advanced for many species (Forchhammer, Post & Stenseth 1998; Parmesan & Yohe 2003; Root *et al*. 2003; Dunn 2004; Thackeray *et al*. 2010). An increase of 1°C has been demonstrated to advance breeding phenology by 3-5 days in a range of species (McLean *et al*. 2016; Phillimore *et al*. 2016; Socolar *et al*. 2017). Functionally, this trend to breed earlier may maintain temporal synchrony with key prey resources (Radchuk *et al*. 2019) which is particularly important for species breeding in highly seasonal environments where prey is ephemerally abundant (Both *et al*. 2009). Asynchrony with a prey peak can negatively impact reproductive output, with negative effects from such trophic asynchrony termed the match-mismatch hypothesis (Both *et al*. 2006; Miller-Rushing *et al*. 2010; Samplonius *et al*. 2021).

The advancement of avian breeding phenology across years and geographic gradients in response to increasing average temperature is well established (Crick *et al*. 1997; Dunn & Winkler 2010), however the impacts of localised intra-annual temperature variation is less understood and limited to use of coarse-grained climatological indices such as North Atlantic Oscillation and average monthly climate variables (Suggitt *et al*. 2011; Huey *et al*. 2012). This is primarily due to a lack of studies collecting fine-scale, within-site territory temperature data alongside bird breeding data, despite this being the scale relevant to individual birds (Scherrer, Schmid & Körner 2011; Hinks *et al*. 2015). Studies show that temperatures within a nest cavity can modify breeding phenology (Simmonds *et al*. 2017), and that fine-scale temperature is a good predictor of localised species distribution (Frey, Hadley & Betts 2016), but it is unknown how territory settlement, breeding phenology and reproductive output are influenced by territory-scale temperature.

Furthermore, in the same breeding locations, resident species with similar ecological requirements to long-distance migrants often commence breeding earlier (Källander *et al*. 2017). With a warming spring climate this difference can become more divergent with resident species more responsive to breeding cues in warm (early) years (Potti 2009; Samplonius *et al*. 2018; Haest, Hüppop & Bairlein 2020), suggesting a potential role of temperature.

Understanding the influence of temperature on territory occupation, breeding phenology and reproductive output could indicate how optimally timed a population is. In the well-studied highly seasonal deciduous forest system, as tree and caterpillar phenology is well synchronised with fine-scale temperature (Buse *et al*. 1999; Van Asch *et al*. 2013), locally warmer territories will show advanced phenology in these trophic levels compared with cooler territories (Cole *et al*. 2015; Hinks *et al*. 2015). Therefore, if breeding consistently commences earlier in warmer territories this could indicate successful thermal niche tracking, where individuals in a well-timed population are able to track the local phenology of lower trophic levels (Dobrowski 2011; Socolar *et al*. 2017). However, if breeding occurs earlier in cooler relative to warmer territories, then the average population consequence could be later than optimal timing as late breeders (either through late arrival, lower quality or competition) are forced to occupy the remaining warmer territories, increasing potential for mismatches with prey resources (Bowers *et al*. 2016). If breeding commences earlier in warmer territories in cool years, and in cooler territories in warm years, average population timing would be optimal in cool years and suboptimal (late) in warm years, although as far as we are aware such hypotheses remain untested.

Examining a population’s breeding phenology across a temperature gradient means few assumptions need to be made about optimal dates or specific prey items compared to studies focussed on dates alone. Optimal synchrony of consumer species breeding in the same locality may vary due to different prey requirements and developmental needs of offspring. For example, breeding tits and flycatchers mostly consume moth caterpillars (Cholewa & Wesolowski 2011; Samplonius *et al*. 2016) and breed at overlapping but different times to one another (Burgess *et al*. 2018; Samplonius *et al*. 2018), so within any single year and locality breeding timing across these species will encompass peaks of different prey species at different nestling developmental stages. Whether breeding is timed to the same or different optima, a preference for breeding in cooler territories first, which have later prey peaks benefiting breeding in warm years when peaks are most advanced, could indicate local population-level mistiming. This territory occupation pattern would also suggest an adjustment of territory selection and breeding phenology in response to the local phenological stage (Hušek, Lampe & Slagsvold 2014; Hinks *et al*. 2015; Shutt *et al*. 2019), in addition to relying on temperature as a causal cue of breeding time (Caro *et al*. 2013; Verhagen *et al*. 2020).

In this study, we investigate how territory-scale temperature variation over four years within a 417ha woodland affects territory settlement, breeding phenology, clutch size and fledging success of three co-occurring, insectivorous woodland passerine birds; long-distance migrants pied flycatcher and wood warbler, and the resident blue tit. We hypothesise that as the two migrants tend to breed slightly later than blue tits, and later breeding in warm years increases asynchrony with prey, that in warm years migratory species will be less optimally timed compared to resident species, selecting cooler territories first and showing lower fledging success in warm territories.

## Methods

This study was conducted at East Dartmoor National Nature Reserve in Devon, South-West England, UK (50.60°N -3.73°E) over four breeding seasons, 2015-2018. Habitat across the reserve is relatively uniform and dominated by sessile oak *Quercus petraea* with a European holly *Ilex aquifolium*, and rowan *Sorbus aucuparia* understory and a European bilberry *Vaccinium myrtillus* field layer. The study area extends linearly approximately 5 km along a steep-sided river valley in the north, with a 150 ha woodland containing two shallow stream valleys in the south, together resulting in slopes at different aspects and an altitudinal range of 60-300 m.

### Breeding bird data

Throughout the study area, 309 wooden nest boxes with a 32 mm entrance hole were available each year and monitored annually throughout the bird breeding season April-June for pied flycatcher and blue tit breeding attempts. Nest box checks were made at least weekly and more frequently at some nest stages to determine species occupancy, date of first nesting material, date of first egg laying, completed clutch size and the number of fledglings for each breeding attempt. Date of first nesting material was taken as the date of the first nest check that found fresh nesting material for nests that ultimately had a clutch of eggs laid in, with nests found half complete (cup shape evident) or more over the maximum 7 days inspection interval assigned the mid-point between the two visits. All species were assumed to lay one egg per day (Perrins 1979; Lundberg & Alatalo 1992), with the first egg date back-calculated from incomplete clutches recorded at nest inspections. Clutch size records the total number of eggs laid that were incubated, and the number fledged was the number alive at the last visit pre-fledging minus the number found dead at the first visit post-fledging. Second broods following successful first broods are rare in these species (Parejo & Danchin 2006; Both, Ubels & Ravussin 2019) and we had no known cases in our study. Although pied flycatchers can be polyterritorial, this is rare in our population with only two cases observed during the study period, both retained in the analysis. Predated nests were included in analyses with few instances in our data for the two hole-nesting species as nest boxes exclude most nest predators.

The date of spring arrival by territorial male pied flycatchers to territories was also recorded from the first week in April. Territorial activity, such as singing, alarm calling and inspection of nest boxes, was observed on alternate mornings, and the presence of, colour and position of any rings and variable plumage characteristics such as forehead patch size were recorded. These were cross-referenced against males captured at the same nest boxes later in the breeding season. Male arrival date to a territory was taken as the mid-point between the date of the survey prior to the first observation of the male and the date of the survey when the male was first observed. This is a common method used for pied flycatchers (Bell *et al*. 2017; Potti *et al*. 2018) with its accuracy verified by geolocator tracking data (Both, Bijlsma & Ouwehand 2016). Nearly all breeding male and female pied flycatchers were captured annually and marked with a unique metal numbered ring upon first capture, enabling the frequency of breeding over an individuals’ lifetime to be determined, including for years prior to 2014 using data collected prior to the present study. Between 2015-2018, 320 pied flycatcher (2015=70, 2016=81, 2017=84, 2018=85) and 315 blue tit (2015=86, 2016=73, 2017=70, 2018=86) nesting attempts were monitored.

Wood warbler territories and nests were also monitored throughout the breeding seasons 2015-2018. Wood warblers are ground nesting and were located first by locating territory holding males from singing behaviour and then by observing parental behaviour in these territories, with the location of each nest determined by GPS and visited every 3-4 days. First egg-laying dates were determined from visits during egg laying assuming that one egg is laid per day. A total of 37 wood warbler nests were monitored during the study period (2015=12, 2016=13, 2017=9, 2018=3). Most breeding male and female wood warblers were captured annually and marked with a unique metal numbered ring and three colour rings upon first capture, which enabled us to determine that there were no cases of second broods following a successful first brood. Although nest predation rates are typically high in wood warblers, in our study six nests were predated at chick stage which were included in analyses.

Territory quality for pied flycatcher and blue tit was inferred from the proportion of years each nest box had been occupied by either nesting pied flycatchers or blue tits out of the total number of years each nest box was available (erected and possible to breed in) since 1980 and until 2018. We pooled the two species as both have similar requirements and occupation by one species in a given year precludes occupation by the other. Long-term nest box or territory occupancy patterns provide a clear correlate of territory quality in other studies, with more frequently occupied territories of higher quality (Sergio 2003; Potti *et al*. 2018). We used the occupancy rate since 1980 as both the number of nest boxes and breeding population size had stabilised by this year after nest box provisioning started incrementally from 1955.

### Temperature data

Between 2015-2018 96 temperature dataloggers (Thermachron iButton DS1921G) were placed throughout the study area at each intersection of a 150 m grid. Each was attached to a tree 1.5 m above ground level in a waterproof standard sized plastic container with three drilled holes to provide air circulation (Shutt *et al*. 2019). Temperature dataloggers recorded temperature hourly to an accuracy of 0.5°C. Additionally, 22 higher-sensitivity temperature dataloggers (Thermachron iButton DS1922L) that record temperature to an accuracy of 0.0625°C were attached alongside 22 of the lower resolution dataloggers on the same trees with the same aspect in a non-random way to maximise the spread of higher-sensitivity loggers across the study area.

We used temperatures recorded between Julian days 97 and 132 (7 April to 12 May in non-leap-years), which encapsulates the time period during which pied flycatcher and blue tit egg laying phenology is most sensitive to thermal variation in the study area (Samplonius *et al*. 2018). Less is known about wood warbler thermal sensitivity periods, but this period corresponds to wood warbler arrival and territory occupation, just before egg laying, so is a feasible time window. Data from temperature dataloggers with more than 16 missing hourly data points within the focal period within a year were discarded as incomplete (n=25, from a total of 333). Additionally, data from temperature dataloggers recording at least one measurement beyond the extreme credible temperature bounds within the focal period (<-8°C or >35°C) were discarded for the year in question as unreliable or faulty (n=9). As all faulty dataloggers were the higher sensitivity model, and data from high sensitivity dataloggers are not directly comparable with data from low sensitivity dataloggers, the unaffected higher resolution datalogger data were only used to determine the accuracy of their paired low sensitivity datalogger and excluded from all other analyses (n=56). Paired high sensitivity and low sensitivity dataloggers gave very similar readings (interquartile range <0.2°C despite recording on different sensitivities with a possible correct error range of 0.25°C), whereas the variation across the entire study area was much higher (interquartile range >0.6°C in each year), providing confidence that local temperature differentials across the study area correspond to genuine local temperature variation rather than datalogger inaccuracy. Temperature means across the focal period were used and a total of 257 datalogger mean readings remained over the four-year period after this quality control (87 in 2015, 54 in 2016, 56 in 2017 and 60 in 2018).

All spatial analysis was conducted within R version 3.6.3 (R Development Core Team 2018) using the raster, gstat and tmap libraries (Pebesma 2004; Hijmans *et al*. 2015; Tennekes 2018). The data from each remaining low sensitivity temperature datalogger within each year was then projected to its geographical location and temperature across the study area interpolated using inverse distance weighting (Pebesma & Heuvelink 2016) raised to the power two. The inverse distance squared weighting power was decided upon after investigating the ability of the data to predict known temperature points in a ‘leave one out’ validation routine of powers between one and four (root-mean-squared-error of P1 = 0.467, P2 = 0.447, P3 = 0.453, P4 = 0.461). Temperature differentials (mean location temperature difference from site yearly mean) for each nest box and wood warbler nest location were then extracted from the resultant heatmap raster (Pebesma & Heuvelink 2016). On inspection, the heatmap differential rasters for each year were highly consistent in both spatial pattern and variance, with cooler locations always cooler and warmer locations always warmer, which was supported by a variance component analysis. Therefore, an interannual differential mean (mean location temperature difference from site yearly mean averaged across all years) was used, which we term the local temperature differential. This allows determination of whether territories were in a relatively cool or warm area within the overall study area across years and afforded greater differential consistency as not every temperature datalogger position recorded data in every year.

### Statistical analyses

All analyses were conducted in R version 3.6.3 (R Development Core Team 2018). General linear models (GLM) were constructed to assess the effects of local temperature differences on measures of reproductive output for this system. Not all breeding parameters were assessed as a response variable for each of the three bird species due to lack of data. Five breeding parameters were used: date of male arrival to breeding territory and date of nest building initiation (both pied flycatcher only), date of first egg laying, clutch size and fledging success (all three species). All dates are reported as April days (days from the 1 April, e.g. 25 April = day 25). For wood warblers, as nests were often found after eggs had been laid, if date of first egg laying was not recorded but date of hatching was, first egg date was inferred by subtracting the mean number of days (n=18) between first egg date and hatching date across those nests where both were recorded, from hatching date where first egg date was not recorded. Wood warbler nests found during the nestling stage (n=15) were not used in the first egg date analysis.

Response variables were tested by Bayesian GLMs in R (Hadfield 2010) with annual mean temperature, local temperature differential, number of years of historical nest box occupancy (as a proxy for territory quality), previous breeding experience and the interaction between annual and local temperature as fixed predictor variables. Continuous predictor variables were mean centred for ease of interpretation and model functioning (Schielzeth 2010). Nest box ID was included as a random effect for pied flycatcher and blue tit models, and male and female ID as random effects for pied flycatcher models. Territory quality and nest box ID could not be included as predictors for wood warbler as this species does not use nest boxes and therefore territories could not be ascribed long-term occupancy rates. Year was not included as a random effect as it covaried with annual mean temperature for each data point, however we ran additional models for each response with year as a random effect but as these did not significantly influence results they are not reported here. Each model was run for 100,000 iterations with the first 10,000 discarded as burn-in with a Gaussian error structure and non-informative parameter-expanded priors, with both response data and model traces visually inspected to ensure the fit was acceptable.

Individual breeding experience was defined as the number of years an individual had previously been captured breeding in the population. Previous breeding experience and parent ID could not be included as predictor variables for blue tit or wood warbler due to incomplete data on parental identity. For pied flycatcher date of arrival, male breeding experience was included as a fixed effect, whereas in all other models female breeding experience was used, as females build the nests, lay the eggs and provide the majority of parental care in pied flycatchers (Lundberg & Alatalo 1992). Pairwise correlation between fixed effects was assessed and as all were <0.18 this suggested no issue with colinearity within the models, with the most significant correlation being that higher quality territories tended to have cooler local temperature differentials.

## Results

Local spring temperature varied by around 2°C from the coldest to warmest areas across the study area with colder areas at the higher elevations of Yarner Wood in the south and Neadon Cleave in the north, and warmer areas at lower elevations along the River Bovey in Lustleigh Cleave (Fig 1). This pattern was spatially consistent across all four years, across years warmer territories were consistently warmer and cooler territories were consistently cooler, with logger ID explaining 89.8% of the total variance of annual differentials in the variance component analysis. Variation in mean spring temperature between years was observed, with 2018 the warmest (mean = 10.7°C) and 2016 the coolest (mean = 9.2°C).

**Figure 1.**
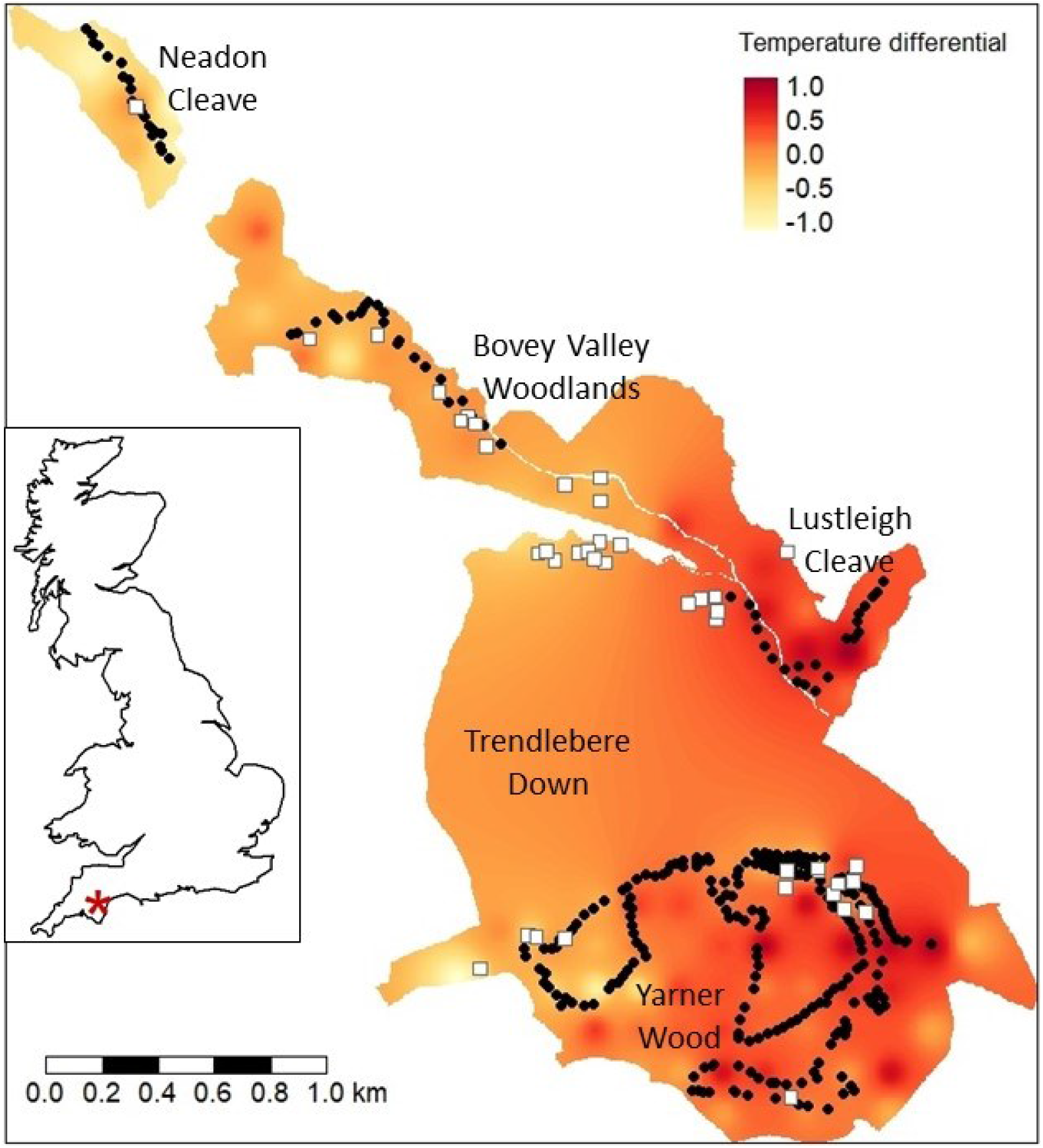
Map of the study area (East Dartmoor National Nature Reserve, Devon, UK), showing its location within Great Britain (inset, red star), the location of nest boxes (black points) and wood warbler nests (white squares), the local names of sections of the reserve, and the relative local temperature differential.

Male pied flycatchers returning from migration in spring occupied the coolest territories first, irrespective of the relative mean annual temperature, with our model predicting arrival to the coolest territories on 12 April and warmest territories on 19 April (Table 1, Fig 2a). There was also an interaction effect of local temperature on the timing of nest building initiation; in cool years nests were initiated significantly earlier at warmer territories, with this pattern reversed in warm years with nests initiated earlier at cooler territories (Table 1, Fig 2b). This temperature effect on the timing of nest building did not translate into an effect on the timing of egg laying or clutch size (Table 1). Fledging success varied by 1.5 fledglings per nest attempt across the temperature gradient, with success higher at the coolest territories compared to the warmest territories (5.6 vs. 4.1 fledglings per attempt respectively) (Table 1, Fig 2c). This latter effect was strongest in cool years, while in warm years fledging success was similar irrespective of territory-level temperature (Table 1, Fig 2d).

**Table 1.** The effects of temperature, territory quality and breeding experience on reproductive phenology and outcomes in three forest passerine species. All results are taken from Bayesian GLMM’s, with slope estimates and credible intervals for each fixed term, with significance asterisks for significant terms (pMCMC ≤0.05* ≤0.01** ≤0.001***), and variance estimates and credible intervals for each random term (the terms in brackets). Yearly temperature and territory quality are mean centred, breeding experience relates to males for arrival date and females for all other measures, and the temperature interaction is between yearly temperature and territory temperature.

| Term | Arrival date | Nest initiation | Egg laying date | Clutch size | Fledging success |
| --- | --- | --- | --- | --- | --- |
| **Pied flycatcher** |  |  |  |  |  |
| Intercept | 15.97 (15.11 – 16.82) | 25.19 (24.26 – 26.16) | 38.26 (37.39 – 39.11) | 6.82 (6.70 – 6.94) | 4.91 (4.60 – 5.23) |
| Yearly temperature | 1.16 (-0.16 – 2.39) | -1.05 (-2.43 – 0.33) | -1.16 (-2.36 – 0.02) | -0.10 (-0.24 – 0.06) | -0.10 (-0.50 – 0.29) |
| Territory temperature | 4.11 (1.75 – 6.49) *** | 0.79 (-1.69 – 3.35) | 0.18 (-1.89 – 2.37) | -0.13 (-0.42 – 0.17) | -0.84 (-1.60 – -0.06) * |
| Temperature interaction | 2.73 (-0.60 – 6.09) | 4.10 (0.21 – 7.93) * | -0.30 (-3.62 – 3.00) | 0.15 (-0.27 – 0.57) | 1.04 (0.00 – 2.19) * |
| Territory quality | -4.77 (-10.40 – 0.74) | -11.22 (-16.81 – -4.76) *** | -6.43 (-11.08 – -1.31) ** | 0.38 (-0.30 – 1.01) | 1.83 (0.06 – 3.64) * |
| Breeding experience | -0.84 (-1.60 – -0.07) * | -0.42 (-1.21 – 0.38) | -0.93 (-1.58 – -0.27) ** | 0.12 (0.03 – 0.22) * | -0.02 (-0.25 – 0.22) |
| (Residual variance) | 20.81 (13.85 – 27.93) | 34.56 (22.96 – 46.52) | 32.21 (23.85 – 40.57) | 0.38 (0.23 – 0.52) | 3.33 (2.13 – 4.54) |
| (Nest box ID variance) | 1.70 (0.00 – 5.80) | 1.35 (0.00 – 4.80) | 1.18 (0.00 – 4.23) | 0.03 (0.00 – 0.11) | 0.39 (0.00 – 1.08) |
| (Male ID variance) | 2.98 (0.00 – 8.57) | 1.45 (0.00 – 5.64) | 1.97 (0.00 – 4.46) | 0.02 (0.00 – 0.07) | 0.72 (0.11 – 1.39) |
| (Female ID variance) | N/A | 12.68 (0.00 – 22.21) | 4.62 (0.00 – 11.03) | 0.33 (0.17 – 0.51) | 0.72 (0.00 – 1.77) |
| **Blue tit** |  |  |  |  |  |
| Intercept |  |  | 24.43 (23.40 – 25.48) | 8.45 (8.20 – 8.69) | 3.10 (2.70 – 3.55) |
| Yearly temperature |  |  | -2.22 (-3.91 – -0.53) ** | 0.20 (-0.16 – 0.60) | 0.78 (0.12 – 1.46) * |
| Territory temperature |  |  | -3.02 (-5.91 – -0.11) * | -0.53 (-1.23 – 0.18) | -0.69 (-1.83 – 0.51) |
| Temperature interaction |  |  | 0.22 (-4.51 – 5.11) | 1.13 (0.09 – 2.18) * | 1.37 (-0.40 – 3.38) |
| Territory quality |  |  | -2.37 (-4.51 – 5.11) | 0.46 (-1.01 – 1.84) | 1.33 (-1.01 – 3.78) |
| (Residual variance) |  |  | 68.09 (56.57 – 80.55) | 2.88 (2.17 – 3.67) | 10.36 (8.36 – 12.56) |
| (Nest box ID variance) |  |  | 2.01 (0.00 – 7.10) | 0.66 (0.00 – 1.30) | 0.88 (0.00 – 2.26) |
| **Wood warbler** |  |  |  |  |  |
| Intercept |  |  | 49.92 (45.12 – 55.08) | 5.59 (5.23 – 5.94) | 2.99 (2.02 – 3.94) |
| Yearly temperature |  |  | 2.57 (-6.54 – 11.40) | 0.35 (-0.28 – 0.97) | 0.09 (-1.56 – 1.89) |
| Territory temperature |  |  | -3.53 (-25.31 – 17.68) | 0.11 (-1.33 – 1.40) | -1.68 (-5.41 – 2.10) |
| Temperature interaction |  |  | 51.59 (13.58 – 91.09) * | -2.82 (-5.23 – -0.50) * | -0.44 (-7.06 – 5.86) |
| (Residual variance) |  |  | 136.2 (59.75 – 237.1) | 1.07 (0.59 – 1.63) | 8.17 (4.57 – 12.39) |

**Figure 2.**
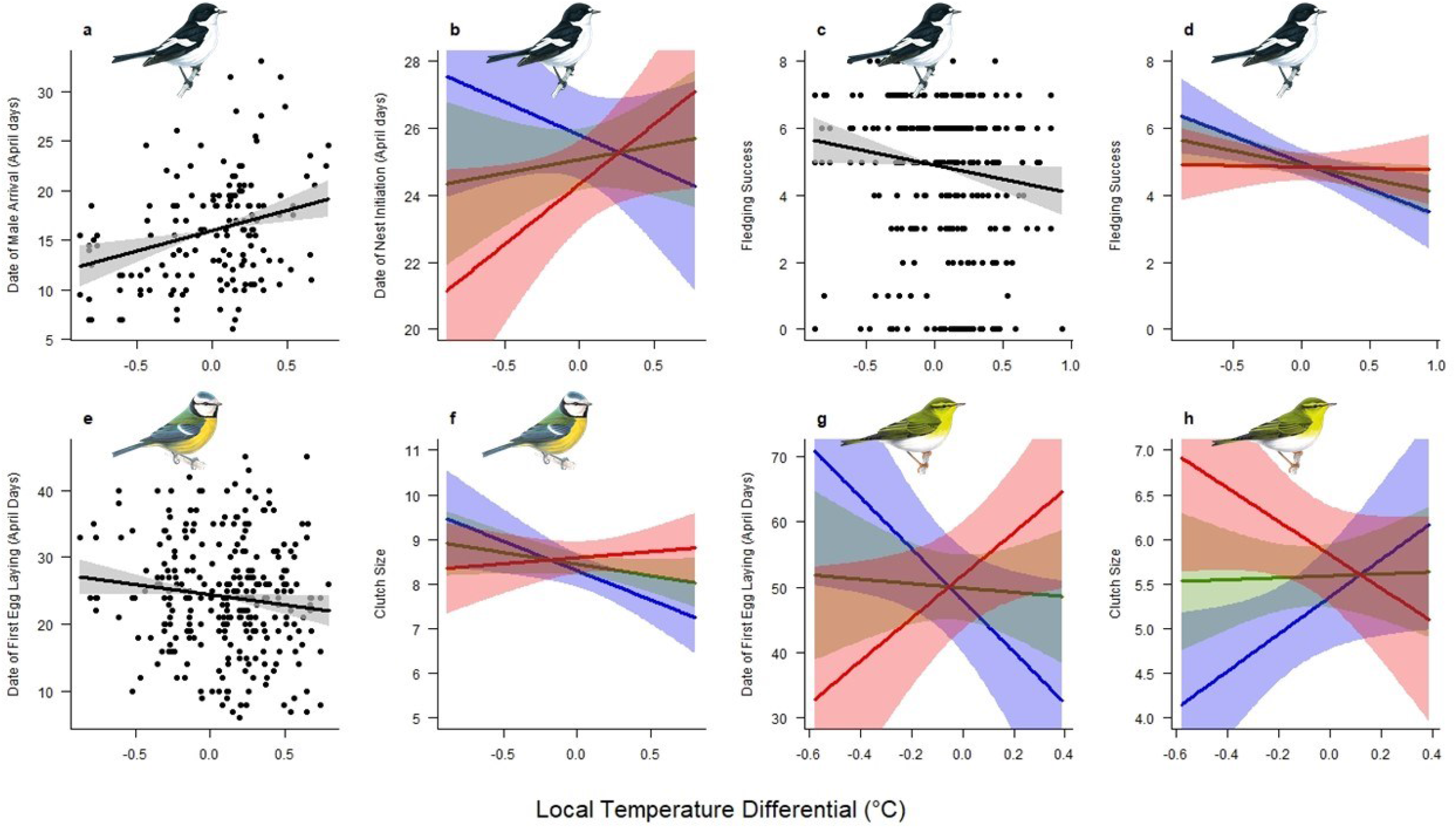
Illustrating the significant effects of territory temperature on the breeding parameters of three woodland passerine species: (a-d) pied flycatcher (e-f) blue tit (g-h) wood warbler. Figures a, c and e show significant single predictor linear effects (solid black line) along with 95% credible intervals (grey shading). Figures b, d and f-h show significant interaction linear effects (solid lines) and 95% confidence intervals (shading) in a cold (blue, -0.7°C below mean interannual temperature), average (green, mean interannual temperature) and warm (red, +0.7°C above mean interannual temperature) year. Bird drawings reproduced with permission from Mike Langman, RSPB Images.

Although the relatively small sample size for the migratory wood warbler means these results had a larger associated uncertainty, there were significant effects of local temperature interactions on wood warbler breeding parameters. In cooler years, eggs were predicted to be laid substantially earlier at warmer territories than cooler territories, whereas in warmer years the opposite was found, with eggs laid earlier at cooler territories (Table 1, Fig 2g). Wood warbler clutches were also significantly larger at cooler territories in warmer years, and at warmer territories in cooler years (Table 1, Fig 2h). There was no significant effect of local temperature on wood warbler fledging success (Table 1).

For resident blue tits, higher local temperatures predicted a significant advancement of five days in the timing of egg laying between the coolest and warmest territories (Table 1, Fig 2e). There was also an interaction effect of local temperature on clutch size, with cooler territories predicted to have substantially larger clutches in cool years, while in warm years warmer territories were predicted to have slightly larger clutches (Table 1, Fig 2f). There was no significant effect of local temperature on blue tit fledging success, although the interaction effect was in the same direction as for clutch size (Table 1). In warm years, blue tits laid eggs earlier (−2.22 days/°C) and had higher fledging success (+0.78 fledglings/°C) (Table 1).

Previous breeding experience (the number of previous breeding seasons individuals had been recorded breeding) also affected pied flycatcher breeding parameters. Males with more experience selected territories earlier (−0.84 days/year experience, Table 1, Fig 3a) and females with more experience laid larger clutches (+0.12 eggs/year experience, Table 1, Fig 3b) earlier in the year (−0.93 days/year experience, Table 1, Fig 3C). Female ID explained more variance than male ID for nest initiation date, egg laying date and clutch size, but they were similar at explaining variance in fledging success. In higher quality territories pied flycatchers also initiated earlier nest building (21 April vs. 1 May, Table 1, Fig 3d) and egg laying (6 May vs. 12 May, Table 1, Fig 3e), and produced more fledglings (4.0 vs. 5.6 fledglings per attempt, Table 1, Fig 3f), with figures at the two extremes of territory quality found. Territory quality showed no significant relationship with blue tit breeding parameters and nest box ID explained little variance across all models (Table 1).

**Figure 3.**
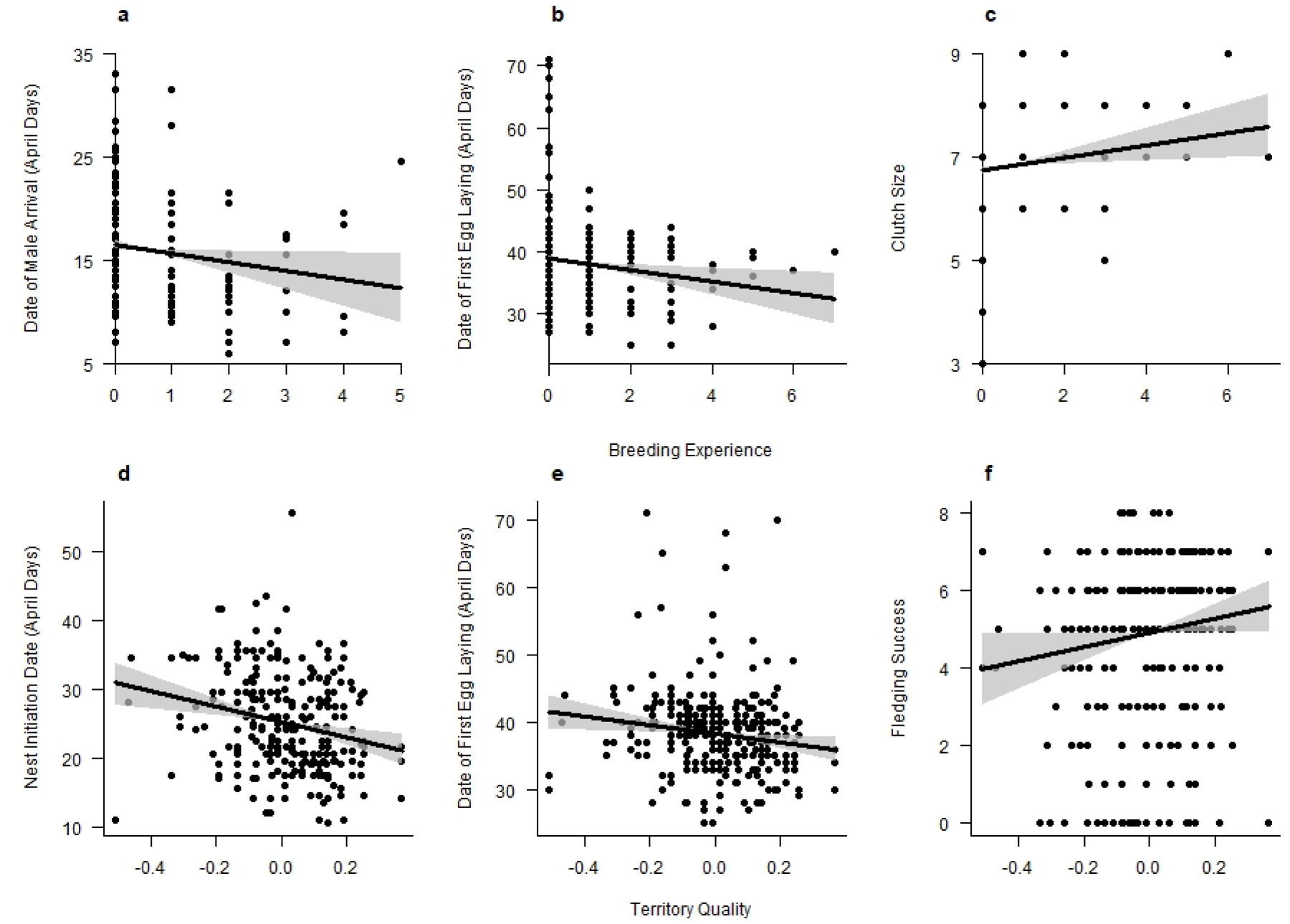
Illustrating the significant effects (solid black lines) of breeding experience (a male and b-c female) and territory quality (d-f) on the breeding parameters of pied flycatcher, along with 95% credible intervals (grey shading).

## Discussion

This study demonstrates multiple relationships resulting from spatial variation in territory-level temperature on territory occupancy, breeding phenology, clutch size and fledging success for three insectivorous woodland birds. Within a 417ha woodland we found both migratory and resident species either selected breeding territories where the temperature would be expected to better synchronise with resource phenology, with these locations differing between cold and warm years, and/or show fitness benefits in relation to the temperature of the territory occupied. In cold years, when vegetation and invertebrate phenology in deciduous woodlands is relatively late, pied flycatcher nest building and wood warbler egg laying commenced earlier in warmer territories, and clutch sizes for wood warbler were larger. In warm years, when vegetation and invertebrate phenology in deciduous woodlands is relatively early, cooler territories were preferred by the migratory species, with earlier nest building by pied flycatchers, and earlier egg laying and larger clutch sizes by wood warblers. Territory-scale temperature also influenced pied flycatcher fledging success across years, with 1.5 more fledglings predicted per nest in the coolest territories compared to the warmest. This pattern was driven by considerably higher fledging success in cool territories in cooler years.

### Temperature impacts on breeding territory settlement and phenology

The role of temperature in animal territory settlement decisions is little explored, with settlement studies including temperature focussing at the scale of distributions (Frey, Hadley & Betts 2016), on the role of environmental factors such as habitat (Jones 2001), on the temperature of nest sites themselves (Hart, Downs & Brown 2016; Carroll *et al*. 2020), or temperature between breeding habitat types (Walsberg 1993; Pollock *et al*. 2015). We expect this paucity of studies is because measuring territory-scale temperature has only recently become achievable with new methods (Haesen *et al*. 2021). Furthermore, while effects of temperature on breeding phenology can be tested experimentally, experimental tests of the role temperatures plays in wild territory settlement is very challenging. Temperature-territory settlement relationships are shown indirectly by an experimental study that manipulated willow warbler *Phylloscopus trochilus* territory settlement, finding female selection of male-held territories was related to male song, and that song rates were higher in warmer territories, which were selected first, and exhibited earlier egg laying (Arvidsson & Neergaard 1991). Also, breeding birds can modulate the sensitivity of hormonal responses to local conditions to fine-tune the onset of territoriality (Wingfield & Hunt 2002).

Pied flycatcher territory and female mate selection is well studied, with males establishing territories and females selecting male-held breeding territories based on habitat quality (Alatalo, Lundberg & Glynn 1986) and male traits (Sirkiä & Laaksonen 2009). Territory choice can be further influenced by information obtained from heterospecifics, for example from cues related to occupancy and density patterns of resident species (Forsman *et al*. 2008) or inspecting multiple nest sites upon arrival (Schuett *et al*. 2017). In our study male pied flycatchers returning from migration settled in cooler territories first. Our measure of arrival was only for males as they are more conspicuous, but our method of estimating nest building initiation is frequently used as a proxy for female arrival date (Potti & Montalvo 1991; Visser *et al*. 2015) as pairing and nest building typically commences one or two days after female arrival (Potti & Montalvo 1991; Lampe & Espmark 2003). If we make that assumption, as nesting begins earlier in cooler territories in warm years and warmer territories in cool years, our results suggest that both male and female pied flycatchers may also partly base territory selection on territory scale temperature variations. Presumably, territories selected first by the earliest arriving birds are those most likely to match with similarly fine-scale variation in vegetation phenology (Hinks *et al*. 2015; Cole & Sheldon 2017) and/or invertebrate prey phenology, and these may mediate avian interactions with temperature. In warmer years migration timings may constrain earlier arrival by migrants (Both & Visser 2001), with the implication that in warm years later arrivals are forced to breed in the warmest territories with a later resource phenology and the greatest potential for mismatch. Therefore, in cooler years, at the local population level, pied flycatchers are likely to be better phenologically matched with resources.

Evidence for a direct relationship between ambient temperature and timing of animal reproduction is mainly demonstrated in birds, and mostly in great tits *Parus major* (Meijer *et al*. 1999; Visser, Holleman & Caro 2009; Verhagen *et al*. 2020), with some evidence of similar relationships in mammals (Kriegsfeld *et al*. 2000; Larkin, Jones & Zucker 2002). Several physiological mechanisms are hypothesised to link temperature with reproduction (Caro *et al*. 2013), and in birds breeding phenology is linked to periods of temperature increase (Schaper *et al*. 2012) and mean pre-breeding temperature (Shutt *et al*. 2019). Also, birds can show stress responses such as an increase in circulating corticosterone in extreme environments or after encountering extreme events such as storms, which could delay breeding onset (Silverin & Wingfield 1998; Wingfield & Hunt 2002).

Egg laying date in our study species is constrained by photoperiod (Lambrechts & Perret 2000; Caro *et al*. 2007), and environmental factors (Shutt *et al*. 2019), and is strongly related to spring temperature at larger spatial scales (Both & Visser 2001; Phillimore *et al*. 2016), with laying starting later at cooler locations such as higher latitudes and elevations (Burgess *et al*. 2018; Bründl *et al*. 2020). We found spatial variation in laying date with regards to territory-scale temperature for both wood warbler and blue tit. Wood warblers started egg laying earlier at warmer territories in cooler years and cool territories in warmer years. Previous work on wood warblers shows a strong relationship between the temperature of the pre-laying period and laying date at larger spatial scales, with higher pre-laying temperatures associated with earlier laying (Wesołowski & Maziarz 2009). We found the same pattern at the territory scale in cool years, however, in warmer years the opposite pattern was exhibited suggesting thermal limits to this pattern. Higher territory temperature linearly predicted earlier egg laying in resident blue tits, similar to the pattern found across longer gradients of space and time but with a slightly shallower slope estimate (McLean *et al*. 2016; Phillimore *et al*. 2016). Our results are consistent with temperature being a direct cue for blue tit breeding phenology (Bourgault *et al*. 2010; Shutt *et al*. 2019) and that this cue applies at a fine scale. We found no significant temperature relationship for pied flycatcher lay date, suggesting temperature may not be a direct cue to egg laying in this species. Pied flycatcher egg laying phenology shows less spatial and inter-annual variation compared with arrival phenology (Nicolau *et al*. 2021), indicating that in years with early arrival individual birds can build nests well in advance of egg laying. This is supported by our results, as in cool years nests were initiated significantly earlier at warmer territories with this pattern reversed in warm years, but these temperature patterns did not significantly impact laying date.

We found that pied flycatcher nest building, and wood warbler egg laying, commenced earlier in warm territories in cool years and in cold territories in warm years. This pattern suggests that breeding timing is optimal in cool years and suboptimal in warm years. Under the match-mismatch hypothesis warmer springs can lead to increased mismatches between prey and food demands of nestlings because phenology is advanced and invertebrate prey peaks are advanced and shorter in duration (Visser *et al*. 1998; Smith *et al*. 2011). Although we did not measure vegetation or prey phenology at the territory scale in our study, the most likely explanation of the observed patterns is territory-scale tracking of resources. The patterns shown in our study also highlight the potential for a plastic response to environmental change by individuals to track breeding resources through adjusting territory settlement decisions (Charmantier & Gienapp 2014).

### Temperature impacts on clutch size

We found spatial variation related to temperature in clutch size for wood warbler and blue tit, but not for pied flycatcher. Wood warbler clutch size followed a similar pattern to breeding phenology, being larger at cool territories in warm years and at warm territories in cool years, with clutch size therefore larger in territories selected first. The effect of territory temperature on clutch size may therefore indirectly contribute to the known seasonal decline in wood warbler clutch size (Wesołowski & Maziarz 2009; Grendelmeier *et al*. 2015), although this could also arise through differences in parental quality in relation to settlement patterns. Although wood warblers do not time breeding to match caterpillar abundance peaks (Maziarz & Wesolowski 2010; Mallord *et al*. 2017), if higher clutch sizes are indicative of better synchrony to other invertebrate prey peaks our results imply that wood warblers may be late matched in warm years and select territories partly based on temperature, refined within territories by topographical and structural nest placement preferences (Pasinelli *et al*. 2016). Blue tit clutch size showed the opposite interaction, with larger clutches in warm territories in warm years and cool territories in cool years. There is evidence that blue tit clutch size has increased with temporal increases in spring temperature, suggesting blue tits can benefit from warmer years (Potti 2009).

Polyterritoriality in well known in pied flycatchers and wood warblers and the second nests of polyterritorial males have later egg laying dates (Temrin 1984; Lundberg & Alatalo 1992) which could have affected our results if this behaviour was common, however we only observed four cases in our study, two pied flycatchers, with differences in associated egg laying dates of 5 and 12 days, and two wood warblers, with differences of 7 days for one with the other unknown due to incomplete data.

### Temperature impacts on fledging success

We found no territory-level temperature effects on fledging success in blue tit or wood warbler, but pied flycatchers showed a strong response, fledging 1.5 more fledglings in cool compared to warm territories, with this effect strongest in cool years. Higher fledging success from cooler territories could be due to these territories having the best match with fine-scale prey phenology, or simply a higher availability of prey. Alternatively the patterns could be explained by physiological relationships between temperature and nestlings that have previously been shown in birds (Andreasson, Nord & Nilsson 2018; Corregidor-Castro & Jones 2021).

### Non-temperature impacts: territory quality and individual experience

For pied flycatchers, our models also controlled for other factors that may influence nest building, clutch size and fledging success. Higher territory quality predicted earlier nest initiation and egg laying, and higher fledging success, supporting the general validity of using the indirect proxy of long-term occupation rates to indicate territory quality in birds (Sergio 2003; Potti *et al*. 2018), particularly when measuring territory quality in relatively homogeneous habitats where habitat composition and structural preferences can be hard to determine or differentiate. The spatial pattern of temperature across the study area was consistent between years, and territories that were relatively cool in 2015-2018 had the highest occupancy rates since 1980, indicating that relatively cooler territories have probably been preferentially selected for almost 40 years. The territory selection relationships we found in our study years probably hold historically as, despite a warming of ambient temperature over these years, breeding phenology at the population level has advanced in response (Nicolau *et al*. 2021), and so the spatial patterns of temperature at the territory scale at the onset of the breeding period remain the same. The models also showed that adult pied flycatchers with greater previous breeding experience (i.e. older individuals) tended to settle on territories earlier in spring (male experience), lay eggs earlier and lay larger clutches (female experience). This is concordant with other studies and is indicative of more experienced parents having higher nesting success (Potti 1998; Both, Bijlsma & Ouwehand 2016; Fay *et al*. 2021).

## Conclusions

Territory-level temperature variations within a single site have profound effects on the breeding decisions and outcomes of all three passerine bird species studied. Resident blue tit breeding phenology responds linearly to increasing territory temperatures similar to responses across large-scale geographic and annual temperature gradients (Phillimore *et al*. 2016). Migrant pied flycatchers and wood warblers, on the other hand, may be responding to small-scale phenological cues at lower trophic levels, and local populations may be late timed in warm years. Current climate change scenarios may therefore potentially favour resident species that are better able to track local phenology across all years (although not always, see Visser *et al*. 1998) over migratory species which may be better synchronised in cool years. Similarly, warm-adapted species respond more positively to current climatic change compared to cold-adapted species (Bowler *et al*. 2015; Scridel *et al*. 2017). As animal distributions are limited by thermal breeding niches (Socolar *et al*. 2017), this contributes to shaping breeding species composition, distribution and abundance (Jiguet *et al*. 2010; Lindström *et al*. 2013). For species conservation aimed at promoting breeding success, knowledge of territory and thermal preferences and their influence on fecundity and fledging success can be used to help inform the location of new forest planting under current large-scale afforestation targets (Burke *et al*. 2021), and conservation interventions such as nest box provision.

